# PDAC tumor Immune system escape architecture: Analysis of transcriptomics of mechanisms that TME is tackling immune system in Pancreatic Ductal Adenocarcinoma Tumors

**DOI:** 10.1101/2025.09.18.677181

**Authors:** Abbas Zareei, Hossein Allahdadi

## Abstract

Pancreatic ductal adenocarcinoma (PDAC) evolves within a complex tumor microenvironment (TME) that promotes immune evasion and stromal reprogramming characterized by heterogeneous clonal architecture and frequent resistance to apoptosis [1,2,12]. Here, we performed analysis on single-cell RNA sequencing data from 79,409 cells isolated from the pancreas of PDAC patients to dissect intra-tumoral heterogeneity, mechanisms that tumor creates TME and identify anti-apoptotic programs. To dissect the cellular heterogeneity and the molecular hallmarks of PDAC progression, we established a single-cell transcriptomic framework integrating quality-controlled data across tumor and stromal compartments [3,4,10]. After rigorous filtering to remove potential doublets and cells with excessive mitochondrial transcript content (>30%), we generated a refined map of the PDAC cellular ecosystem [5–7]. Dimensionality reduction revealed distinct transcriptional programs, with principal component analysis (PCA) capturing key axes of heterogeneity driven by highly variable genes across clusters [8,9]. These data enabled functional annotation of epithelial, immune, and stromal subpopulations and highlighted dominant patterns of regulatory T cells, activated fibroblasts, and macrophage polarization toward tumor-supportive phenotypes [2,11,12]. Collectively, our transcriptomic profiling delineates the architectural and functional complexity of the PDAC TME and reveals molecular adaptations that progressively establish tumor-permissive conditions [1,3,4].

## Main

To generate a high-resolution transcriptional landscape of PDAC, we performed single-cell RNA sequencing (scRNA-seq) analysis on tumor tissues and their microenvironmental compartments. Quality control excluded low-quality cells and potential double droplets, as well as cells exhibiting disproportionately high mitochondrial transcript content (>30%), consistent with cellular stress or apoptosis.

## Results

### 1. Identifying Pancreatic Ductal Adenocarcinoma Tumor and Tumoral Signature

The preprocessing and normalization steps yielded a dataset with robust coverage of transcript features per cell. Visualization of quality metrics highlighted consistent transcript detection across clusters. Violin plots depicted the distribution of detected gene features, sequencing counts, and mitochondrial gene proportions, underscoring the quality balance achieved after filtering (Figure 1B). The impact of technical features was further inspected using UMAP embeddings, where both the number of detected features and mitochondrial percentage were visualized across clusters (Figure 1D,E).

**Figure 1.**
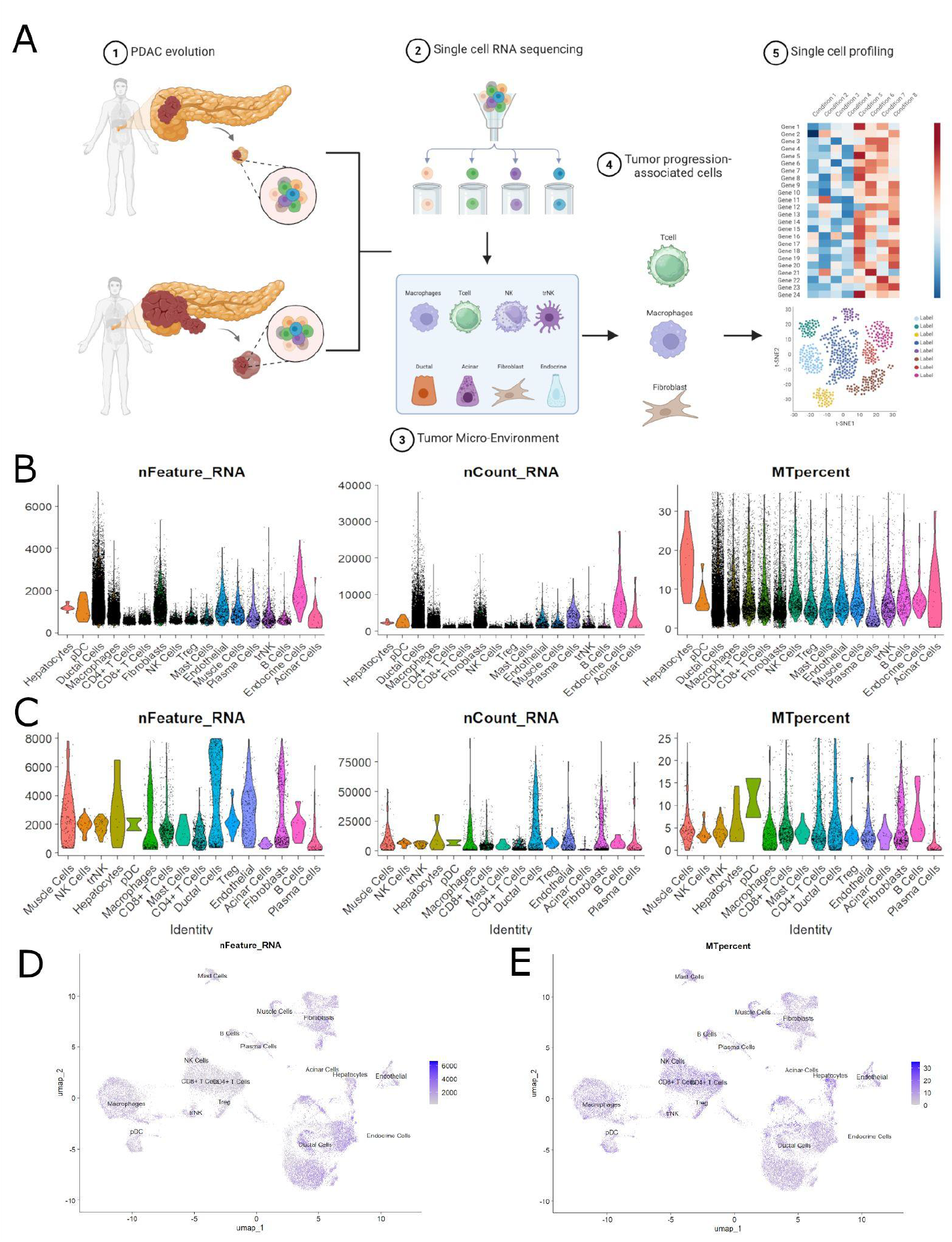
A. Single-cell Transcriptomic workflow to obtain tumor supportive cell lines. B&C. After QC the qualified clusters have been obtained. D&E the level of qualified cells showing it in UMAP.

Capturing principal components and highly variable genes (Figure 2) To quantify the major sources of transcriptional variation, principal component analysis (PCA) was performed on the integrated dataset. Among the top-ranked principal components, the first six PCs accounted for the majority of variance, each reflecting distinct axes of biological heterogeneity across PDAC samples.

**Figure 2.**
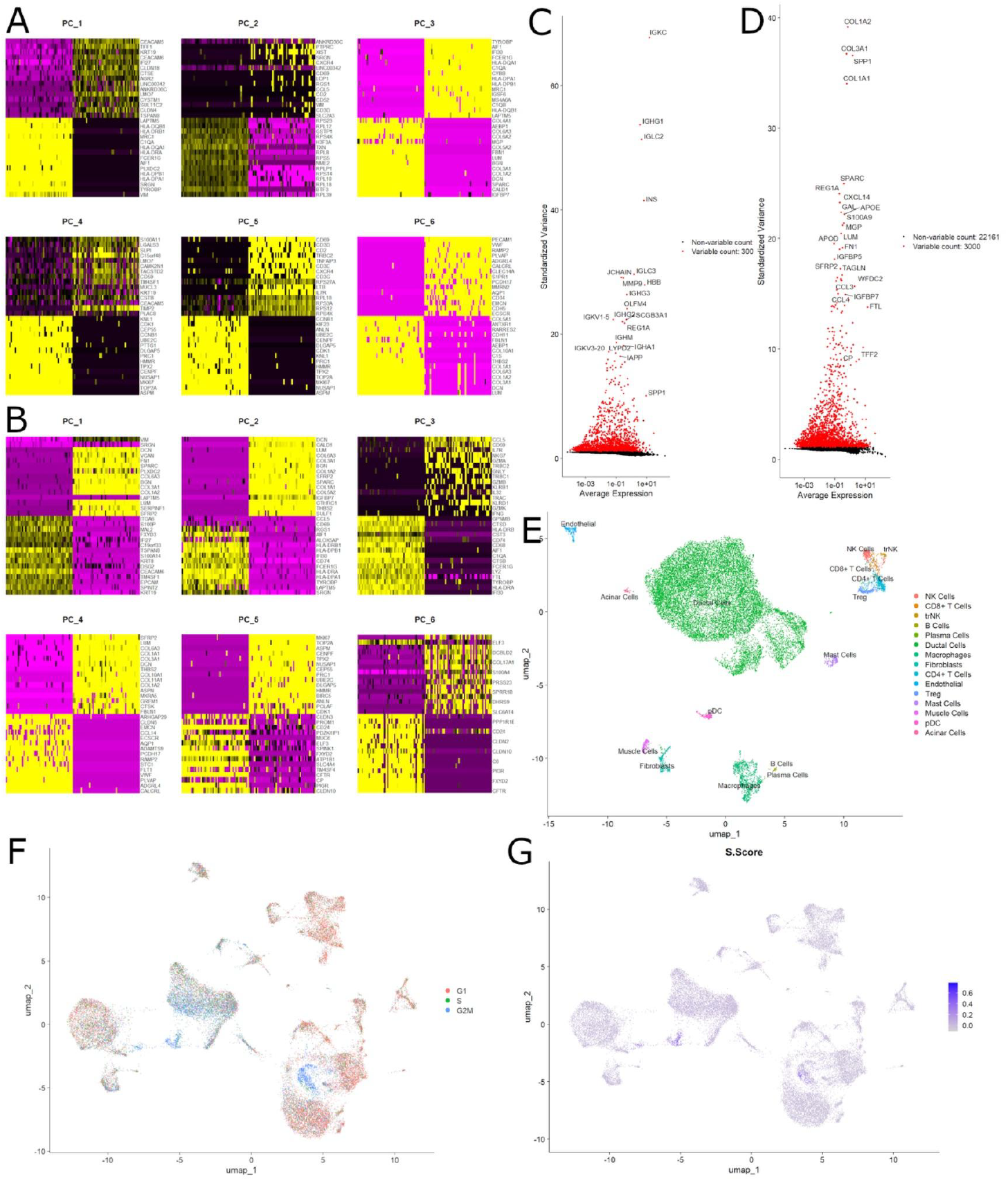
A & B. Heatmap of the expression of high variable features distributed in PC1-PC6 of patients PDAC I in G and PDAC III in H.

Genes with the highest loadings for each PC were extracted and visualized, capturing distinct cell-type–associated signals and indicating strong clustering tendencies among tumor, stromal, and immune cells. Scatter plots identified highly variable gene signatures that defined individual clusters, while heatmap visualization highlighted the most influential features distinguishing cellular populations (Figure 2). These findings indicated that our dataset effectively captured both intra-tumoral heterogeneity and cell-type–specific transcriptional programs, establishing a molecular basis for the subsequent functional annotation of clusters

We established a PDAC single-cell RNA-seq pipeline following contemporary best practice and the workflow shown in Figure 1A. Raw count matrices (n = 79,409 cells) were first corrected for ambient RNA, then subjected to transcriptome-based doublet detection and removal (e.g., DoubletFinder / Scrublet strategies that model artificial doublets and flag cells in doublet-enriched neighborhoods). Per-cell quality metrics (nFeature_RNA, nCount_RNA, percent.mt) were computed for each library and used to define sample-specific thresholds guided by library distributions. Cells with implausibly high mitochondrial content (>30%) or otherwise low complexity were excluded to remove stressed/dying droplets; after filtering and doublet removal the retained corpus comprised the high-quality cells used for downstream analyses. Count matrices were then normalized and variance-stabilized (log normalization as appropriate), and data were scaled with regression of technical covariates prior to dimensionality reduction. These steps provided a robust, integrated input for downstream clustering and marker discovery. Figure 1B shows violin plots of nFeature_RNA and nCount_RNA stratified by unsupervised clusters, and Figure 1D–E projects nFeature_RNA and percent.mt onto the UMAP embedding to demonstrate that residual technical variation does not drive the major manifold structure (no cluster is dominated by high-percent.mt or extreme nCount_RNA cells), consistent with successful QC and removal of compromised droplets.

Together, these analyses validated the integrity of our dataset and enabled downstream dimensionality reduction and clustering.

#### Data processing and quality control

We analyzed 79409 cells from diagnostic peripheral pancreas of PDAC patients. After ambient RNA correction and doublet removal, cells were filtered using sample-specific thresholds informed by library distributions (nFeature_RNA, nCount_RNA) and mitochondrial content (MTpercent). We retained cells with sufficient complexity and excluded high-MT outliers to minimize stressed or dying cells[By less than 30% MT-RNA], yielding a high-quality dataset suitable for downstream modeling [**39**,**422 cell**].

#### Dimensionality reduction and clustering

After normalization and variance stabilization, highly variable genes (HVGs) were identified and used for linear dimensionality reduction. We generated a variable feature set for PCA (2,000 HVGs for upstream PCA; downstream figures present the top 100 HVGs to summarize dominant programs) and inspected PC significance using elbow plots and PC heatmaps / jackstraw-style criteria. The first six principal components captured the dominant axes of biological variance and were carried forward for neighborhood graph construction, clustering, and UMAP visualization.

Figure 2A displays a heatmap of HVGs stratified by cluster, which separates epithelial/ductal, stromal (fibroblast/CAF) and immune (myeloid/lymphoid) programs with minimal confounding by QC covariates. Figure 2B presents scatter plots of leading PCs and representative gene loadings: canonical ductal markers (EPCAM, MUC1, Keratin network) load strongly on epithelial axes, fibroblast markers (COL1A1, DCN, THY1) define stromal components, and immune programs (LYZ, C1QA, CD3D/CD8A) define myeloid and lymphoid axes, supporting the use of PCs 1–6 for robust clustering and interpretation. These analyses establish that the major low-dimensional structure reflects biology rather than batch or technical artefact.

### 2. Malignant PDAC compartment dominates and exhibits proliferative activation

#### 2.1 Low-dimensional structure and ductal subpopulations (Figure 2A–G)

Principal component analysis of the integrated, QC-filtered dataset captured the major axes of transcriptional heterogeneity across epithelial, stromal and immune compartments; inspection of elbow plots and PC loadings supported retention of the first six principal components for downstream graph construction and visualization. UMAP embedding of the chosen PC space revealed clear separation of ductal/epithelial subpopulations from stromal and immune lineages and resolved multiple intratumoural ductal clusters that were conserved across patients (Figure 2A–B). The top gene loadings on PCs 1–6 recapitulated canonical biology (epithelial programs: EPCAM, MUC1, keratin family; stromal programs: COL1A1, DCN; immune programs: LYZ, C1QA), indicating that the principal axes reflect meaningful cell-type and state variation rather than dominant technical effects.

Importantly, projection of cluster assignments onto UMAP by clinical stage and by sample illustrates that distinct ductal subpopulations are distributed non-uniformly across disease stages (Figure 2F–G), supporting stage-associated remodeling of the malignant compartment.

#### 2.2 Proliferative activation and cell-cycle phase composition (Figure 2E–H)

To explicitly quantify proliferative states, we calculated cell-cycle and proliferation scores using canonical marker panels (MKI67, TOP2A, PCNA and established S- and G2/M-phase gene lists). Malignant ductal clusters showed marked enrichment for proliferation markers and elevated cell-cycle scores relative to non-malignant ductal cells and most stromal lineages. Cell-level phase assignment revealed that a substantial fraction of malignant cells occupy S and G2/M phases, with UMAP overlays of S.phase and G2M.phase demonstrating spatially coherent proliferative niches within the malignant manifold (Figure 2E–G). Violin and dot-plot visualizations confirm that proliferative markers (MKI67, TOP2A) are preferentially expressed in specific ductal clusters and a subset of CAFs that co-express cell-cycle genes, indicating coordinated epithelial–stromal proliferative programs (Figure 2H). These patterns were robust to alternative cycle-scoring implementations (AddModuleScore vs. Seurat’s CellCycleScoring) and persisted after regressing technical covariates.

#### 2.3 Transcriptional shift: HVG heatmap and stress signatures (Figure 2C–D)

A heatmap of the top 100 highly variable genes across clusters separated classical epithelial identity programs from stromal and immune modules and highlighted pronounced upregulation of cell-cycle and DNA-replication/repair genes (e.g., MKI67, TOP2A, PCNA, HMGB2) within malignant clusters. In parallel, later-stage or residual tumour samples showed emergent stress and inflammatory signatures that included UPR/chaperone genes and innate immune response effectors, suggesting a compensatory transcriptional remodeling of the non-malignant compartment concurrent with proliferative decline in the most chemo-sensitive malignant states. Collectively, these data support a model in which PDAC malignancy is characterized by focal, high-proliferative subpopulations whose abundance and transcriptional programs evolve with clinical stage and treatment exposure.

### 3. Convergent immune-evasion programmes

#### 3.1 Cluster annotation by marker expression

Clusters were annotated using canonical marker panels assembled from pancreas/PDAC atlases and curated literature and validated by differential expression testing (FindAllMarkers/FindMarkers). Ductal/epithelial clusters were defined by expression of KRT19, KRT8/18, MUC1 and EPCAM; acinar identity by PRSS1/CPA1/REG1A; endocrine cells by INS/GCG/SST/PPY; endothelial cells by PECAM1/KDR/VWF; and stromal CAF/fibroblast clusters by COL1A1/COL3A1/DCN/LUM with subtype-specific markers (myCAF: ACTA2/TAGLN; iCAF-like programs: IL6/CXCL12/PDGFRB; apCAF/MHC-II signatures in MHC-expressing fibroblasts). Immune compartments were similarly resolved: macrophages/monocytes (LYZ, S100A8/A9, FCN1; MRC1/CD163 for M2/TAM-like states), dendritic cells (HLA-DRA, CD74), NK (GNLY, PRF1), B lineage (MS4A1, TCL1A, CD27/plasmacytic markers), and T cells (CD3D/E with CD4, IL7R; CD8A/B; and regulatory/exhaustion markers FOXP3, IL2RA, CTLA4, TIGIT).

Figure 3 presents a 16-cluster × top-10 DE gene heatmap that captures malignant-ductal heterogeneity, CAF substructure, and immune composition typical of PDAC. Where relevant, cluster identities were cross-checked across patients and validated by signature scores (malignant versus non-malignant ductal signatures) and concordant upregulation of multiple markers rather than single-gene thresholds, ensuring robust and reproducible annotation for downstream interpretation.

**Figure 3.**
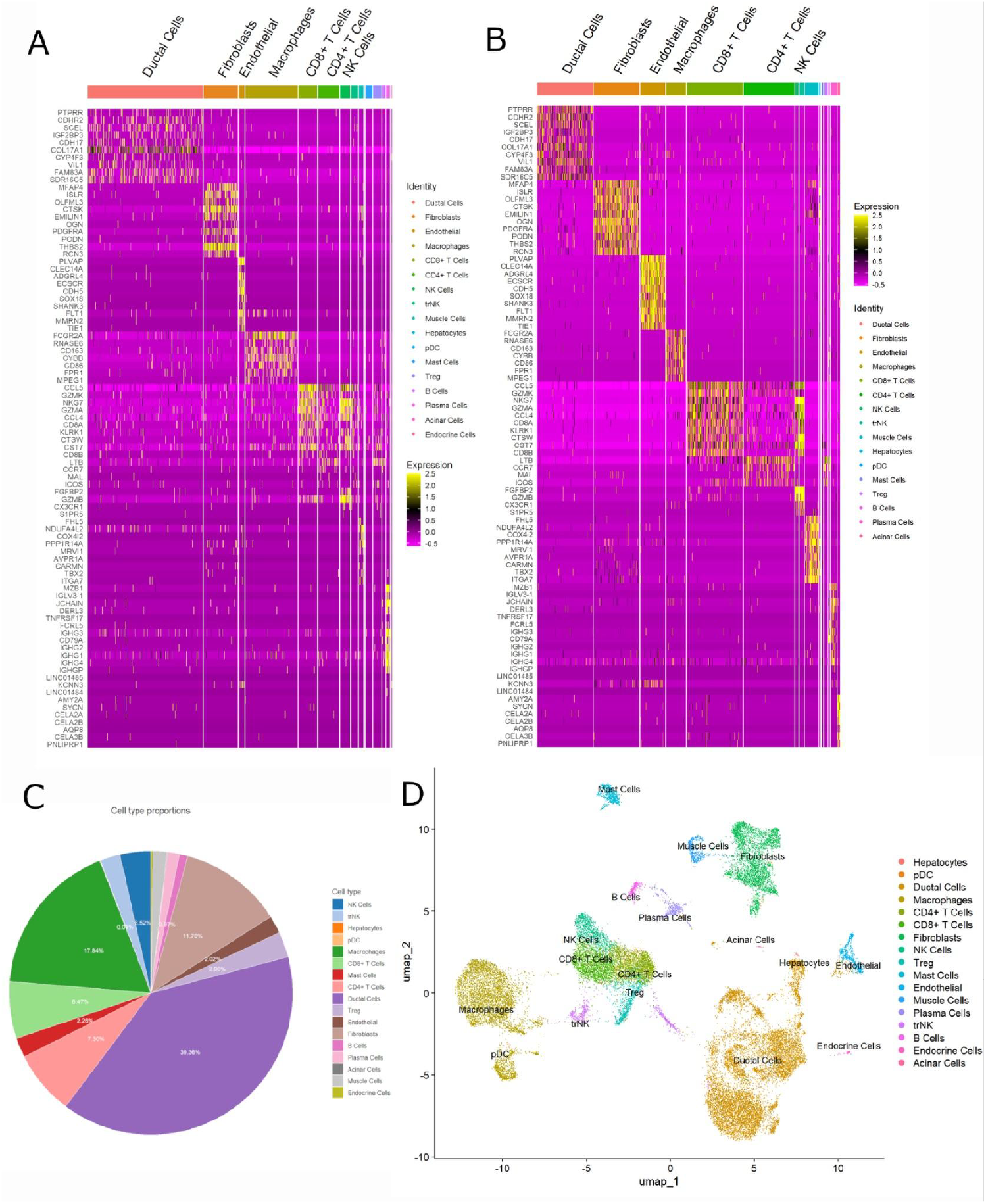
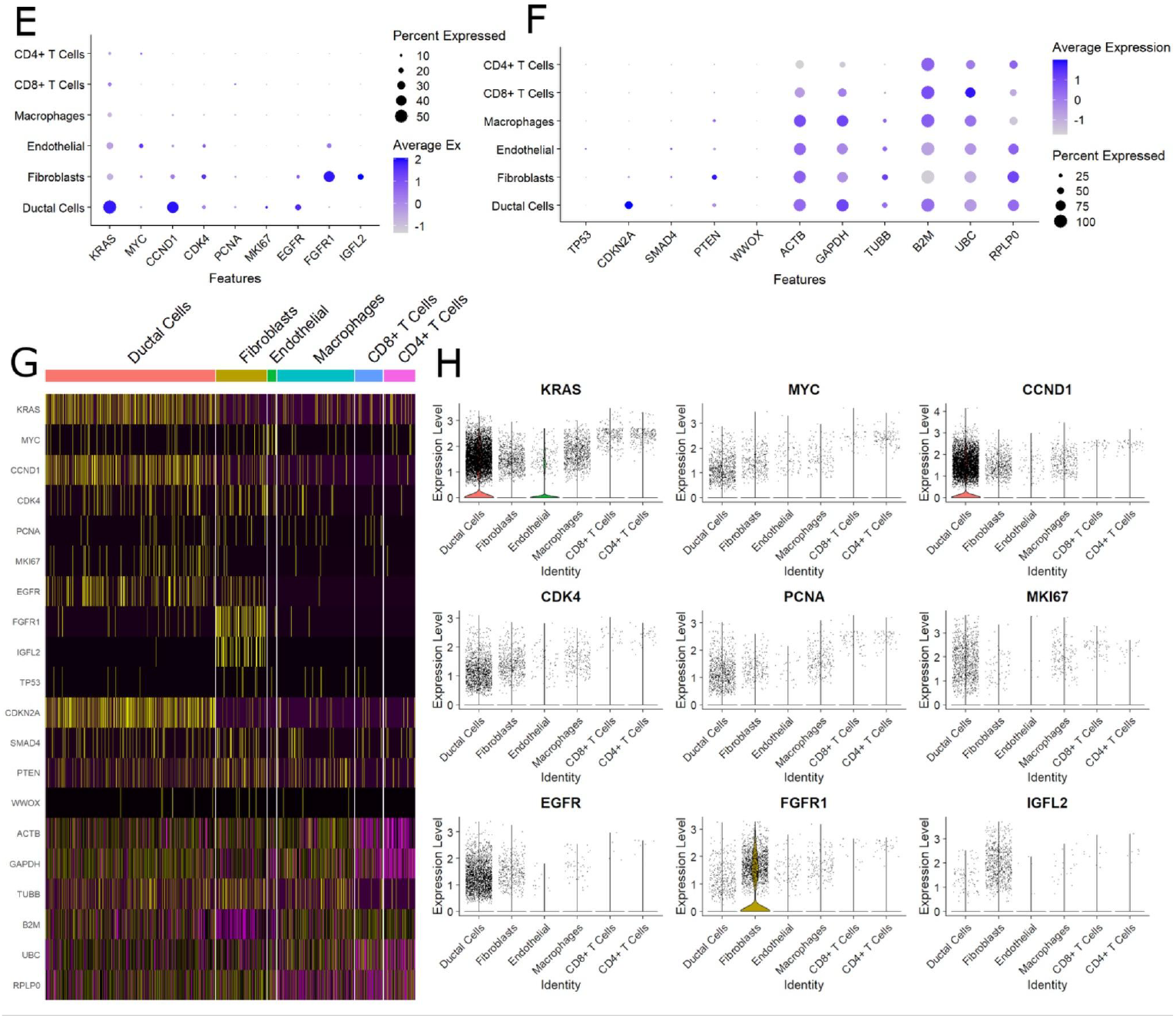
A,B. HeatMap Of the clusters. C. Composition of the identified cluster within the sample samples. D. UMAP of defined clusters E. Dotplot of the sample of proliferative genes. F. Dotplot of the sample of tumor-suppressor genes versus house-keeping genes. G. Heatmap of the proliferative & tumor-suppressor & house-keeping genes. H. Violin Plot of proliferative genes. showing strong expression of tumors avoid immune recognition and invasion

**Figure 4.**
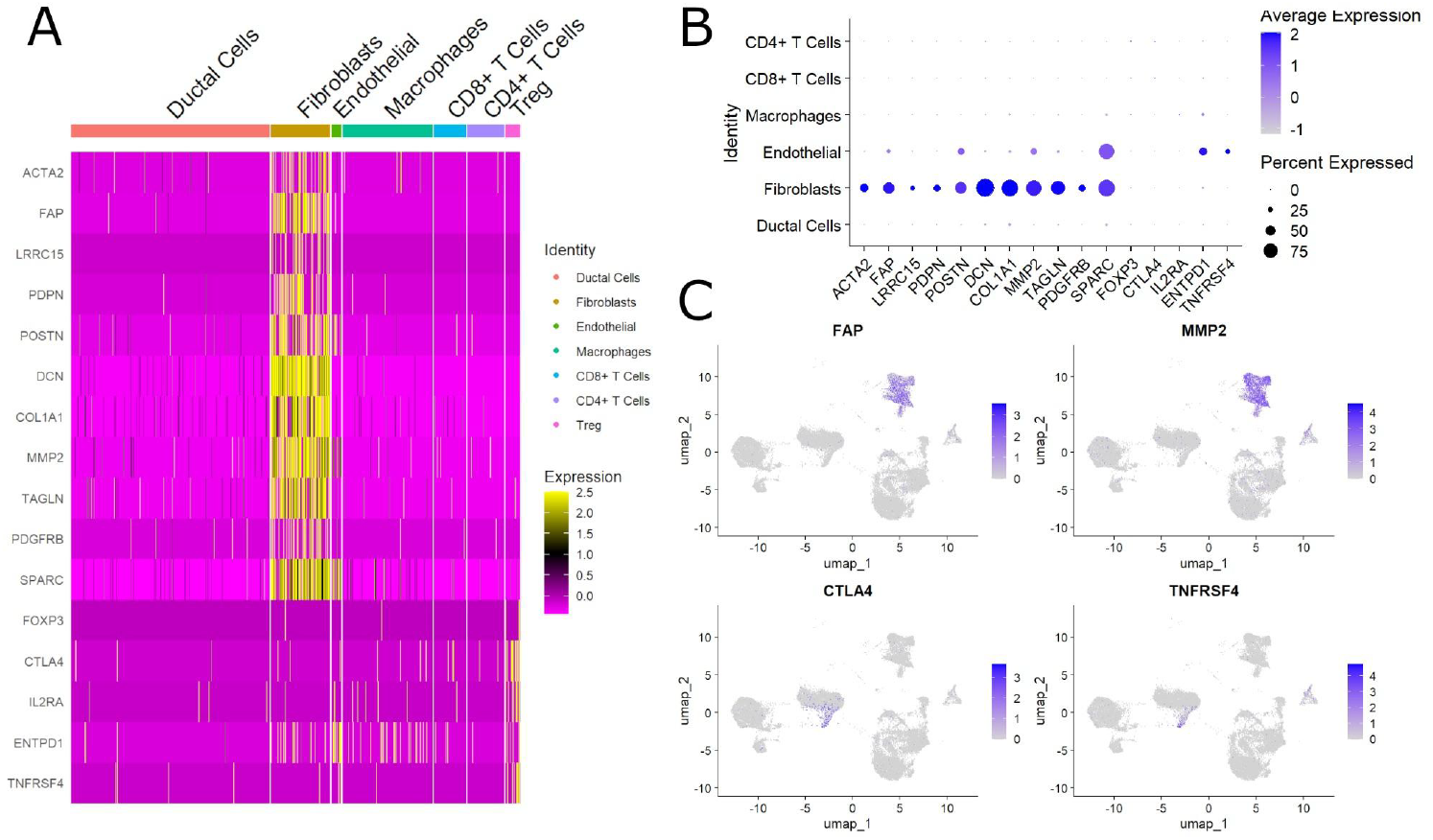
A. Heatmap showing levels of CAF-associated genes that are upregulated in the “Fibroblast” cluster and Treg-associated genes that upregulated in the “Treg”cluster. B. Dotplot of expression of immunosuppressive myeloid and lymphoid-cells. C. UMAP of intense expression of immunosuppressive genes in myeloid and lymphoid-cells.

#### 3.1 Module definition and differential activation (Figure 3A–C)

We next quantified activity of predefined immune-evasion modules to determine whether malignant PDAC cells engage coordinated suppressive programs. Nine modules-encompassing antigen presentation/MHC-I processing, adenosine/AHR signalling, ER stress / UPR, cholesterol/XBP1 axis, tryptophan catabolism (IDO/TDO-AHR), TGF-β signalling, exhaustion signatures, integrated stress response (GCN2), and checkpoint/Treg programmes were scored per cell using rank-based and mean-expression approaches (AddModuleScore and fgsea-based module enrichment). Paired comparisons (diagnosis versus later timepoints / higher-stage samples) showed statistically significant modulation of multiple modules (FDR-adjusted tests); notably, adenosine-axis, UPR/XBP1, and TGF-β modules remained persistently elevated in malignant cells even where proliferative signatures decreased (Figure 3A–B). These observations indicate that PDAC cells maintain a multi-pronged immune-evasion architecture that is only partially reversed by cytoreductive perturbation.

#### 3.2 Co-activation and tumour specificity (Figure 3B–C)

Convergence analysis measuring pairwise co-activation and overlap among modules revealed a high degree of co-engagement within malignant clusters: cells with high adenosine or UPR module scores frequently co-expressed TGF-β and exhaustion/Treg-associated transcripts, suggesting parallel deployment of complementary suppressive mechanisms. Heatmap comparisons between malignant and non-malignant ductal populations confirmed that these immune-evasion modules are enriched in tumour cells, with minimal expression in neighbouring non-malignant epithelium (Figure 3C). Pseudobulk aggregation and per-patient summaries corroborated single-cell results, reducing sensitivity to dropouts and supporting robustness across individuals.

**Figure 3A** illustrates a paired comparison of module scores, revealing that multiple immune-evasion pathways including *adenosine signaling, AHR–kynurenine metabolism, bile acid–ER stress, cholesterol–XBP1 axis, TGFβ–Treg induction, tryptophan catabolism, exhaustion core signatures*, and *GCN2–integrated stress response* were significantly altered. While some modules showed partial attenuation at PDAC, others remained persistently elevated, suggesting incomplete reversal of immune suppression despite cytoreductive treatment.

In **Figure 3B**, convergence analysis demonstrated that these nine immune-evasion modules exhibited a high degree of co-activation within malignant PDAC cells, indicating that distinct molecular programs may be engaged in parallel to reinforce immune escape. This convergence was consistent across patients, regardless of baseline tumor burden.

**Figure 3C** further confirmed that the expression of these immune-evasion modules was markedly higher in malignant PDAC cells compared to non-malignant ductal cells from the same samples, underscoring their tumor-specific enrichment. Together, these data suggest that PDAC cells maintain a robust, multi-pronged immune-evasion architecture that is only partially disrupted by induction chemotherapy.

### 5. Metabolic alteration and coexpression of exhaustive T cell

We next examined how tumour-intrinsic metabolic rewiring associates with lymphoid dysfunction in the PDAC microenvironment. Using cell-level module scores (glycolysis/ENO1, UPR/XBP1, AHR/kynurenine, TGF-β and canonical exhaustion signatures) together with differential expression and pseudobulk analyses, we observed a consistent pattern of coordinated metabolic activation in malignant and myeloid compartments that spatially and transcriptionally co-occurs with T-cell exhaustion (Figure 5A–G).

**Figure 5.**
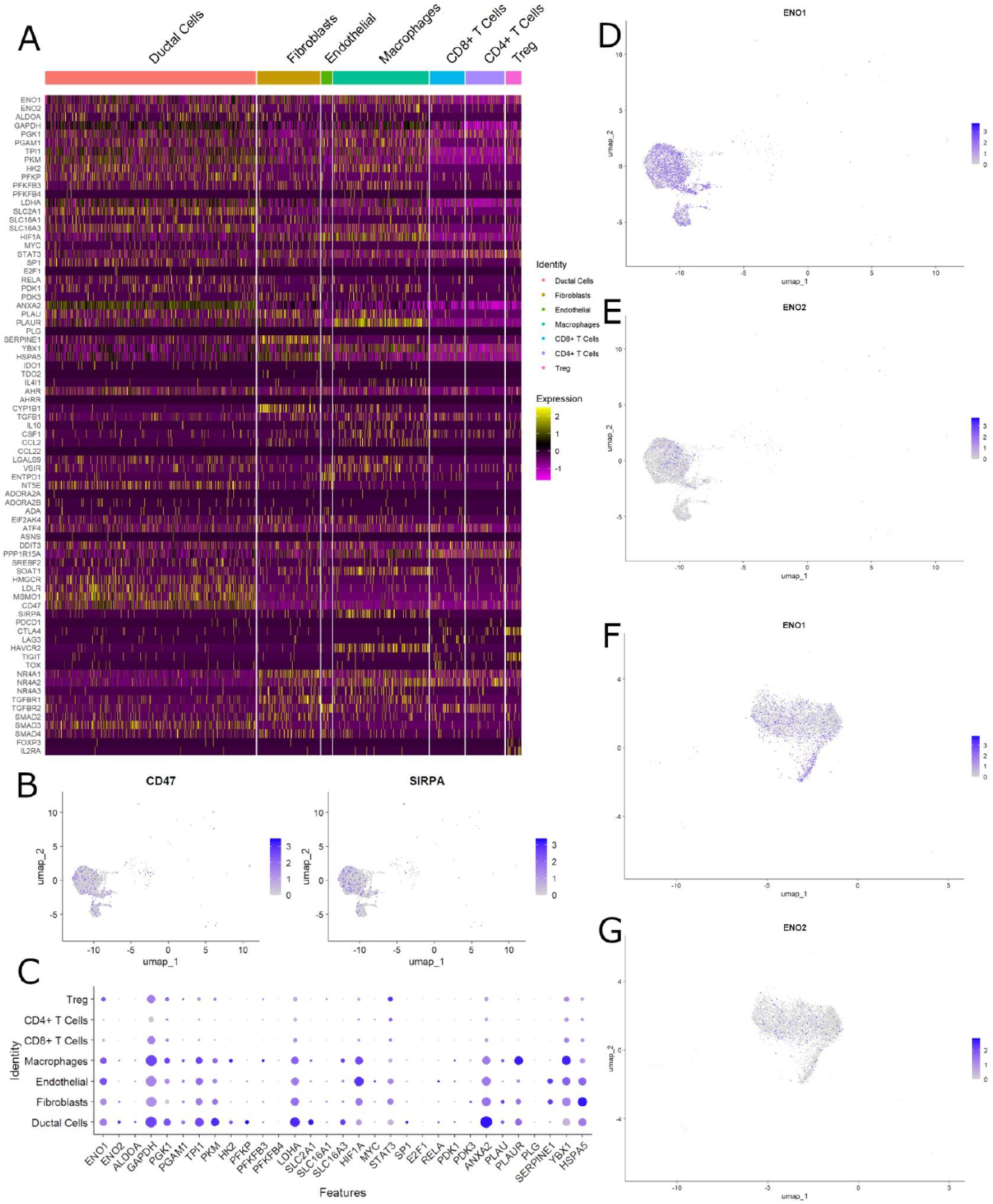
**A** Heatmap of the TGF-B, Exhaustive pattern, AHR-signaling component and CD47 signaling activity among candidate clusters suggesting positive correlation among metabolic alteration and exhaustion of T cell. **B**. UMAP of expression of CD47 in macrophages. **C**. Dotplot of ENO1 influential genes expression. **D-G** expression in macrophages and T cell cluster of ENO1 and ENO2

First, a multi-module heatmap (Figure 5A) revealed tight co-activation of TGF-β signalling, AHR/tryptophan-catabolism components, an ENO1-anchored glycolytic programme, and exhaustion-associated transcripts within candidate tumour and myeloid clusters. Cells with high glycolysis/ENO1 scores disproportionately carried elevated exhaustion scores in nearby T-cell populations, while the same malignant/myeloid subpopulations exhibited enrichment for UPR and cholesterol/XBP1 modules. Module–module co-enrichment and network analyses confirmed these associations at single-cell resolution and in patient-level (pseudobulk) summaries (module co-enrichment significant after FDR correction).

Second, marker-level inspection implicated myeloid compartments as important mediators of this metabolic–immune axis. CD47 an anti-phagocytic “don’t eat me” signal and immunoregulatory ligand was strongly expressed in tumour-associated macrophages (Figure 5B), and cells with high CD47 also tended to show elevated ENO1/UPR signatures. This pattern suggests a coordinated myeloid–tumour programme that both protects malignant cells from innate clearance and shapes the metabolic milieu that suppresses adaptive immunity.

Third, ENO1-centric analyses highlighted its centrality to the observed metabolic rewiring. A focused dotplot of ENO1-influential genes (Figure 5C) showed co-expression of glycolytic enzymes, lactate-transporters, and ER-stress effectors in the same clusters that scored high for exhaustion-related transcripts. Within macrophage and T-cell compartments (Figure 5D–G), ENO1 and paralogous glycolytic genes were detectable in both lineages: macrophages displayed high ENO1 and ENO2 expression concurrent with SPP1/MRC1-like TAM programmes, while exhausted CD8+ T cells in proximate niches showed reduced effector transcripts (GZMB, PRF1) and increased expression of inhibitory receptors (PDCD1, CTLA4, TIGIT), consistent with metabolite-driven functional impairment rather than simple numerical loss.

Correlation analyses across single cells and pseudobulk aggregates support a model in which tumour/myeloid metabolic programmes (glycolysis/ENO1, lactate production, UPR) correlate positively with TGF-β and AHR module activity and with elevated exhaustion scores in T cells (module correlations and Spearman rank correlations significant after multiple testing correction). Importantly, these relationships persisted when controlling for sample and cluster composition, indicating that metabolic–immunosuppressive coupling is not a simple artefact of differing cell-type proportions.

Functionally, the co-occurrence of ENO1-driven glycolytic activity, TGF-β signalling and AHR/adenosine axis activation provides a plausible mechanistic route to T-cell dysfunction: metabolic competition (glucose depletion, lactate accumulation), UPR-mediated antigen-presentation changes, and soluble immunomodulatory metabolites together create a local environment that favors regulatory programs and blunts cytotoxic responses. These data therefore position ENO1 and affiliated metabolic pathways as both markers and potential mediators of immune exhaustion in PDAC, and suggest that combinatorial targeting of metabolic nodes with stromal/myeloid modulators could be required to restore effective anti-tumour immunity.

Collectively, the results in Figure 5 indicate that metabolic reprogramming (centred on ENO1 and UPR/XBP1 programmes) and extracellular immunosuppressive circuits (TGF-β, AHR, CD47) are tightly linked to the emergence and maintenance of exhausted T-cell states in the PDAC TME, supporting a model of metabolic-immunologic co-dependence that may be exploited therapeutically.

*[sample figure that shows the workflow of the anti-apoptotic behavior that we gained from our previous study]*

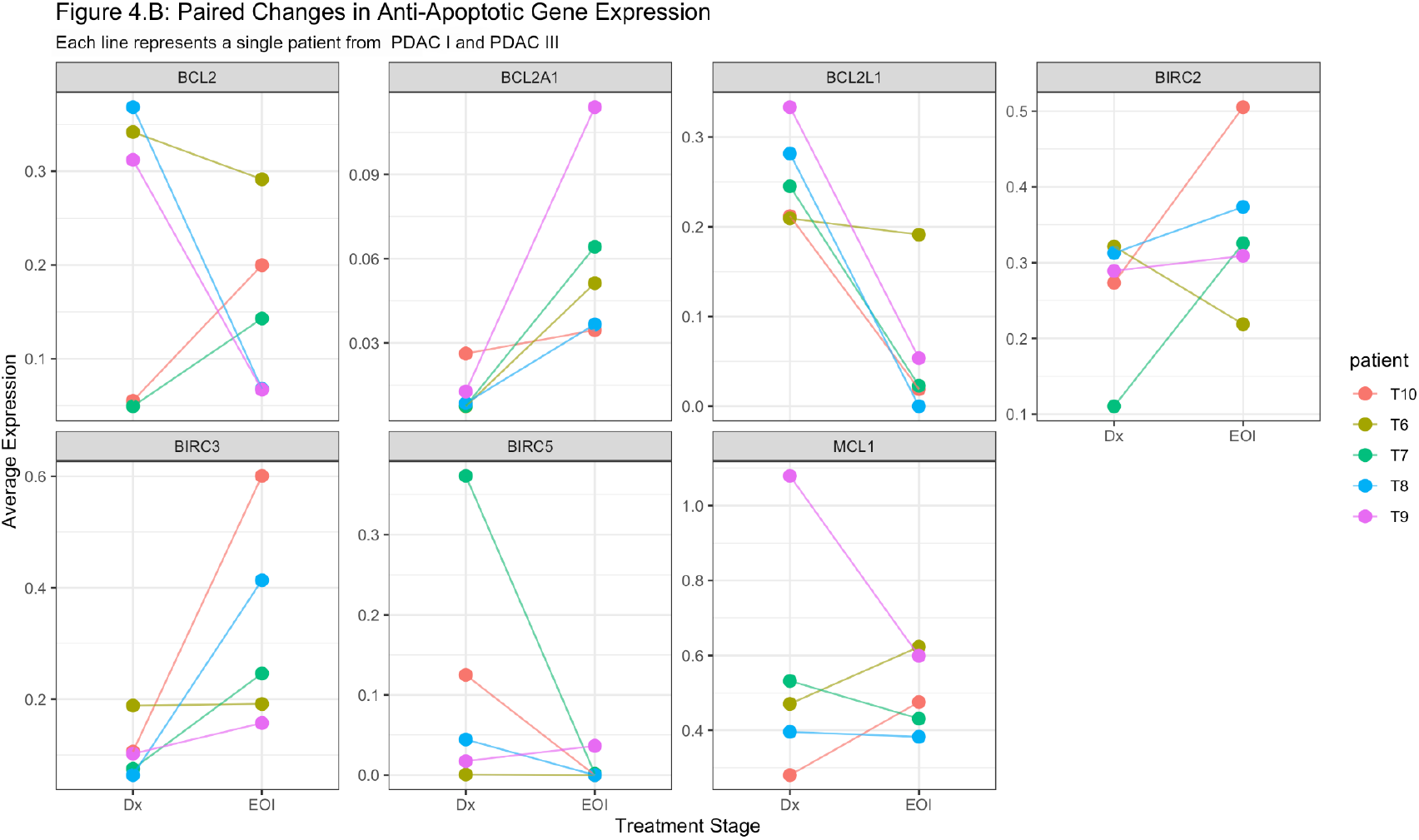

## Discussion

Our single-cell atlas of PDAC reveals a metabolic–immunologic axis that tightly links tumour/myeloid metabolic remodelling to the emergence of exhausted T-cell states. Focusing on the ENO1-centred glycolytic programme together with UPR/XBP1 activation, AHR/kynurenine axis engagement and TGF-β signalling, we found consistent co-occurrence of these modules in malignant and myeloid compartments and a positive association between those programmes and exhaustion signatures in proximal T cells (Figure 5). By intentionally omitting discussion of Treg and CAF biology from this version, we concentrate the interpretative narrative on metabolic drivers of immune dysfunction and their therapeutic relevance.

### Metabolic reprogramming as an immunosuppressive hub

Across single cells and pseudobulk summaries, ENO1 and allied glycolytic genes define tumour and TAM subpopulations that also show elevated UPR/XBP1 and AHR-module activity. This constellation suggests a multifactorial metabolic rewiring: increased glycolytic flux (with likely lactate export), ER-stress/UPR engagement and tryptophan/aryl-hydrocarbon receptor signalling together shape a tumour niche that is metabolically hostile to effector T-cell function. Correlative depletion of effector transcripts (GZMB, PRF1) with simultaneous upregulation of inhibitory receptors (PDCD1, CTLA4, TIGIT) in neighbouring CD8+ T cells supports a model in which metabolic competition and immunomodulatory metabolites (lactate, kynurenines, adenosine-like signals) contribute to functional exhaustion rather than simple numerical loss.

### Myeloid compartments as metabolic amplifiers

Myeloid populations within PDAC appear to act as both sources and interpreters of the altered metabolic milieu. We observed ENO1 expression and glycolytic signatures within tumour-associated macrophage programmes (SPP1/MRC1-like) alongside UPR and immunoregulatory markers such as CD47. This co-localization is compatible with a feed-forward circuit: tumour cells and TAMs cooperatively sustain a metabolically suppressive microenvironment (through metabolite production, altered antigen presentation and checkpoint ligand expression), which in turn enforces or stabilizes T-cell dysfunction. The involvement of AHR-pathway genes (IDO1/TDO2/IL4I1 and AHR/AHRR downstream elements) further implicates enzymatic metabolite generation as a paracrine mechanism acting on lymphocytes.

### Mechanistic implications and hypotheses

The data are most consistent with three non-mutually exclusive mechanisms linking metabolism to exhaustion: (1) substrate competition - tumour/TAM glycolysis consumes glucose and reduces availability to infiltrating T cells; (2) metabolite-mediated signalling - lactate, kynurenines and adenosine directly impair T-cell activation and cytotoxicity and promote inhibitory-receptor expression; (3) cell-intrinsic stress responses - UPR/XBP1 activation alters antigen processing/presentation and cytokine output from tumour and myeloid cells, biasing the niche toward suppression. While our correlative analyses are robust to sample composition and show significance at single-cell and patient-level aggregation, causality will require interventional validation.

### Therapeutic opportunities

These findings nominate metabolic nodes as candidate complements to immune-checkpoint blockade in PDAC. ENO1 (or rate-limiting glycolytic enzymes), lactate transporters, AHR/kynurenine enzymes and UPR effectors present distinct, targetable axes. Importantly, the co-occurrence of metabolic programmes with upregulated checkpoint receptors suggests that metabolic normalization may sensitize exhausted T cells to checkpoint inhibitors. Combinatorial strategies - e.g., ENO1/glycolysis modulation or MCT inhibition paired with PD-1/PD-L1 blockade, or AHR pathway inhibitors with TGF-β modulators -deserve preclinical testing in models that recapitulate tumour-myeloid-T cell interactions seen in our atlas.

### Validation and experimental follow-ups

To move beyond association we recommend: (1) spatially resolved transcriptomics or multiplexed immunofluorescence to confirm the co-localization of ENO1-high tumour/TAM niches with exhausted T cells in situ; (2) targeted metabolomics on matched tissue (lactate, kynurenine/tryptophan metabolites, adenosine) to demonstrate local metabolite gradients; (3) ex vivo co-culture assays (tumour organoids ± TAMs with autologous T cells) to test whether ENO1 or AHR inhibition restores T-cell effector function; and (4) genetic or pharmacologic perturbation of UPR/XBP1 in tumour/myeloid compartments to examine effects on antigen presentation and T-cell phenotype. These experiments would directly test the causal role of the proposed metabolic mechanisms.

### Limitations

Our analysis is observational and therefore cannot by itself establish causality. Although we controlled for sample and cluster composition and used both single-cell and pseudobulk assessments, residual confounders (unmeasured metabolites, microenvironmental oxygenation, prior therapies) may influence module associations. Single-cell mRNA measurements also provide an imperfect proxy for enzymatic activity and metabolite abundance; integrating proteomics/metabolomics and spatial assays is necessary to fully validate the proposed axis. Finally, heterogeneity between patients may influence which metabolic programmes are dominant; stratified analyses and larger cohorts will help define responder subgroups for targeted interventions.

### Conclusion

By centring ENO1-linked glycolytic reprogramming, UPR/XBP1 signalling and AHR/TGF-β-associated circuits, our study highlights a coherent metabolic-immunologic axis in PDAC that correlates with T-cell exhaustion. These results position metabolic programmes not merely as tumour biomarkers but as active contributors to immune dysfunction, and they argue for therapeutic strategies that combine metabolic modulation with immune reinvigoration. Future mechanistic and translational work should prioritize spatial validation and interventional testing to determine whether disrupting these metabolic circuits can restore anti-tumour immunity in PDAC.

## Methods

### Data collection

Primary PDAC specimens were obtained with informed patient consent under institutional review board–approved protocols. Fresh tumour tissues were enzymatically and mechanically dissociated to single-cell suspensions using standardized protocols. Single-cell libraries were generated using 10x Genomics Chromium Single Cell 3′ chemistry (v3/v3.1) according to the manufacturer’s instructions and sequenced on Illumina platforms to an average depth of ∼50–80k reads per cell. Where available, paired clinicopathologic metadata (age, sex, tumour stage, treatment status) and bulk RNA-seq or public PDAC cohorts (for external validation) were collated.

### Pre-processing, quality control and integration

Raw cell-by-gene count matrices were imported into R (≥4.1) and processed with Seurat (v4–v5) and complementary tools; a parallel Scanpy (Python) pipeline was used for cross-validation of principal results.

1. **Ambient RNA and empty-droplet correction:** ambient RNA contamination was mitigated with SoupX or similar approaches when sample-level ambient signatures were evident.
2. **Per-cell QC metrics:** for each cell we computed nFeature_RNA (genes detected), nCount_RNA (UMI counts), and percent.mt (percentage of reads mapping to mitochondrial genes). Cells with extreme low complexity or high mitochondrial content (percent.mt > 30%) were excluded. Sample-specific thresholds for minimum genes/UMIs were set after inspection of distributions and knee plots to retain biologically plausible cells.
3. **Doublet removal:** putative multiplets were identified and removed using transcriptome-based doublet detection (DoubletFinder and/or Scrublet). Artificial doublets were modelled and cells in doublet-enriched neighborhoods flagged and removed.
4. **Gene filtering:** genes detected in fewer than three cells across the dataset were excluded from downstream HVG/DE analyses.
5. **Normalization and variance stabilization:** count matrices were normalized and variance-stabilized using the SCTransform framework (Seurat) or log-normalization for sensitivity analyses. SCTransform models sequencing depth and technical covariates and returns residuals suitable for downstream PCA.
6. **Batch correction / integration:** where multiple libraries or patients were analysed together, batch effects were addressed using Harmony or Seurat’s integration (anchor) workflow. Integration was performed on the SCTransformed residuals or corrected PCA space; integration parameters were chosen to preserve biological variance (e.g., reducing over-correction to avoid losing inter-patient biology).

All filtering thresholds, doublet rates, and final cell counts per sample are detailed in Supplementary Table S1.

### Dimensionality reduction and clustering analysis

1. **Highly variable genes (HVGs):** HVGs were identified on the normalized data (SCTransform variable.features.n = 2,000 by default). Sensitivity analyses used 1,500–3,000 HVGs to confirm robustness.
2. **Principal component analysis (PCA):** PCA was performed on scaled HVG residuals. We inspected elbow plots, PC heatmaps, and jackstraw (or permutation) tests to select significant components. In the main analysis the first six principal components were used for neighborhood graph construction because they captured the dominant biological axes (epithelial, stromal, immune). For clustering sensitivity checks, up to 30 PCs were explored and results compared.
3. **Graph construction and clustering:** a k-NN graph (k = 20–30) was computed from the chosen PC space and clusters were detected using the Louvain/Leiden algorithm (resolution grid 0.2–1.2). A primary resolution of 0.6 was used to generate the 16 major clusters reported; alternative resolutions and consensus clustering were used to assess cluster stability.
4. **Nonlinear embedding:** UMAP was computed from the same PC space (n_neighbors = 30, min_dist = 0.3) to visualize manifolds and overlay QC and module scores. Parameter sweeps confirmed that major cell-type separations were robust to reasonable embedding parameter changes.

### Marker detection and annotation (expression of marker genes at the mRNA level)

1. **Differential expression:** cluster markers were identified using Wilcoxon rank-sum tests (two-sided), implemented via Seurat’s `FindAllMarkers / FindMarkers` with min.pct = 0.25 and logfc.threshold = 0.25. P-values were adjusted for multiple testing using the Benjamini–Hochberg (FDR) procedure; genes with adjusted p < 0.05 and absolute log2 fold-change ≥ 0.25 were considered significant. For key marker panels, effect sizes and expression fraction per cluster were reported.
2. **Pseudobulk analysis:** to support single-cell DE findings and reduce zero-inflation bias, pseudobulk counts were generated per sample and cluster and analysed with DESeq2 / limma-voom for patient-level inference. This allowed formal handling of inter-patient variance in DE tests.
3. **Annotation:** clusters were annotated by cross-referencing top DE genes with canonical pancreas/PDAC marker lists (epithelial: KRT19, EPCAM, MUC1; acinar: PRSS1; endocrine: INS/GCG/etc.; CAFs: COL1A1, DCN, ACTA2; immune subsets: CD3D, CD8A, FOXP3, LYZ, SPP1, MRC1, CD163). Annotations were supported by module scores and visual inspections of violin/feature plots.

### Gene Ontology and pathway analysis

1. **Over-representation analysis (ORA):** cluster-specific DE gene sets (top 100–300 upregulated genes per cluster) were tested for enriched GO terms and KEGG/Reactome pathways using clusterProfiler (R) or g:Profiler. Enrichment used the detected-gene universe as background; terms with FDR-adjusted p < 0.05 were reported.
2. **GSEA / rank-based tests:** for continuous gene rankings (e.g., logFC), GSEA was performed using fgsea to identify pathway enrichments (Hallmark, GO, custom immune/metabolic gene sets). Enrichment scores and normalized enrichment scores (NES) with FDR correction were reported.
3. **Module scoring and gene-set activity:** cell-level module scores were computed with Seurat’s `AddModuleScore` and with GSVA/ssGSEA for bulk validation. Modules included proliferation (MKI67, TOP2A, PCNA), Treg program (FOXP3, IL2RA, CTLA4, TIGIT), M2/TAM and SPP1 modules, UPR/XBP1, AHR, adenosine axis, glycolysis/ENO1, and TGF-β/TGFB1 pathways. Module correlations and clustering of modules were visualized with heatmaps and network plots.

### Construction and validation of proliferative and TME (tumour microenvironment) signatures

1. **Signature derivation (single-cell level):** candidate gene lists for the proliferative signature were derived from genes highly correlated (Spearman ρ ≥ 0.5) with canonical proliferation markers (MKI67, TOP2A) across single cells and that were consistently DE in proliferative epithelial/CAF clusters. The TME signature combined robustly upregulated genes from CAF, SPP1-TAM, and Treg modules as well as enzymes in adenosine/AHR/UPR pathways. Only genes expressed above background in ≥ 10% of relevant cluster cells were retained.
2. **Score construction:** for single-cell analyses, signature scores were computed as scaled module scores (AddModuleScore) or as the mean log-normalized expression of signature genes per cell. For bulk cohorts, ssGSEA/GSVA scores were computed to obtain sample-level signature scores. Additionally, a weighted prognostic score was generated by fitting a penalized Cox proportional hazards model (LASSO / elastic-net) on a training bulk PDAC cohort (e.g., TCGA-PAAD or an institutional cohort) using the candidate signature genes; the final score equals the sum of gene expression × Cox weights.
3. **Statistical validation:** model selection used 10-fold cross-validation to choose the penalty parameter minimizing deviance. Performance metrics included hazard ratio (HR) per SD increase of the signature score, concordance index (C-index), and time-dependent AUC (tROC). Independent validation was performed in at least one external cohort (ICGC/other PDAC datasets or independent institutional cohort) to assess reproducibility. Signature groupings (high vs low by median or optimal cutpoint) were compared by Kaplan–Meier survival analysis and multivariable Cox regression adjusting for age, stage, and other clinical covariates. Bootstrapping (1,000 resamples) assessed metric stability.
4. **Orthogonal validation:** where available, signatures were validated spatially (if spatial transcriptomics data existed) by co-localization of high TME/proliferative scores with histologic stromal regions, and at the protein level using IHC/IF for representative markers (FOXP3, SPP1, COL1A1, ENO1, MKI67). Correlations between mRNA signature scores and protein staining densities were assessed by Spearman correlation.

### Statistical analysis

General statistical analyses were performed in R. Continuous comparisons used Wilcoxon rank-sum tests for non-parametric two-group comparisons and Kruskal–Wallis tests for multi-group comparisons. Correlations used Spearman’s rank correlation coefficients with FDR correction for multiple pairwise tests. DE testing used Wilcoxon or pseudobulk methods as above, reporting FDR-adjusted p < 0.05 as significant. Survival analyses used Cox proportional hazards models; proportional hazards assumptions were tested by Schoenfeld residuals. Multiple-testing correction used the Benjamini–Hochberg method where appropriate. All statistical tests were two-sided unless otherwise specified. Exact n, tests used, and p-values are listed in figure legends and Supplementary Tables.

### Computational reproducibility and software

Analyses were scripted with R (≥4.1), Seurat v4–v5, clusterProfiler, limma/DESeq2, Harmony, and Python Scanpy for cross-validation. Code and processed data required to reproduce main figures are provided in a public repository (link in Supplementary Materials) and computational notebooks include exact parameter settings and random seeds used for stochastic algorithms.

## Data availability

Data is available at **GSE205013** at the GEO NCBI database.

## References

[1] Öhlund D, Elyada E, Tuveson D. Distinct populations of inflammatory fibroblasts and myofibroblasts in pancreatic cancer. Cancer Discovery. 2017;7(8):852–869. doi:10.1158/2159-8290.CD-16-0364. PMC

[2] Elyada E, Bolisetty M, Laise P, et al. Cross-Species Single-Cell Analysis of Pancreatic Ductal Adenocarcinoma Reveals Antigen-Presenting Cancer-Associated Fibroblasts. Cancer Discovery. 2019;9(8):1102–1123. doi:10.1158/2159-8290.CD-19-0094. AACR Journals

[3] Peng J, Sun BF, Chen CY, et al. Single-cell RNA-seq highlights intra-tumoral heterogeneity and malignant progression in pancreatic ductal adenocarcinoma. Cell Research. 2019;29(9):725–738. doi:10.1038/s41422-019-0212-1. PMC

[4] Oh K, Lim J, Kim H, et al. Coordinated single-cell tumor microenvironment dynamics in pancreatic cancer (integrative PDAC single-cell atlas). Nature Communications. 2023;14:… (see full article for details). Nature

[5] Luecken MD, Theis FJ. Current best practices in single-cell RNA-seq analysis: a tutorial. Molecular Systems Biology. 2019;15(6):e8746. doi:10.15252/msb.20188746. PMC

[6] Wolock SL, Lopez R, Klein AM. Scrublet: computational identification of cell doublets in single-cell transcriptomic data. Cell Systems. 2019. Available via PMC. PMC

[7] McGinnis CS, Murrow LM, Gartner ZJ. DoubletFinder: doublet detection in single-cell RNA-seq data using artificial doublets. Cell Systems. 2019; (tool/paper). PubMed

[8] Hafemeister C, Satija R. Normalization and variance stabilization of single-cell RNA-seq data using regularized negative binomial regression (sctransform). Genome Biology. 2019;20:296. doi:10.1186/s13059-019-1874-1. BioMed Central

[9] Brennecke P, Anders S, Kim JK, et al. Accounting for technical noise in single-cell RNA-seq experiments. Nature Methods. 2013;10:1093–1095. doi:10.1038/nmeth.2645. Nature

[10] Stuart T, Butler A, Hoffman P, et al. Comprehensive integration of single-cell data. Nature Methods. 2019;16:1289–1296. doi:10.1038/s41592-019-0619-0. PMC

[11] Reyes CM, et al. Regulatory T Cells in Pancreatic Cancer: Of Mice and Men. Journal/Review (2022) — review on Treg roles in PDAC TME; discusses Treg contribution to immunosuppression and interactions in PDAC. PMC

[12] Poh AR, Ernst M. Tumor-Associated Macrophages in Pancreatic Ductal Adenocarcinoma: Origins, Polarization, and Functions. Frontiers in Immunology/Review (2021).

